# The integration of large-scale public data and network analysis uncovers molecular characteristics of psoriasis

**DOI:** 10.1101/2021.05.10.443441

**Authors:** Antonio Federico, Alisa Pavel, Lena Moebus, David McKean, Giusy del Giudice, Vittorio Fortino, Catherine Smith, Stephan Weidinger, Emanuele de Rinaldis, Dario Greco

**Author notes:** Correspondence to Dario Greco.

## Abstract

In recent years, a growing interest in the characterization of the molecular basis of psoriasis has been observed. However, despite the availability of a large amount of molecular data, many pathogenic mechanisms of psoriasis are still poorly understood. In this study, we performed an integrated analysis of 23 public transcriptomic datasets encompassing both lesional and uninvolved skin samples from psoriasis patients. We defined comprehensive gene co-expression network models of psoriatic lesions and uninvolved skin. Moreover, we collected, curated and exploited a wide range of functional information from multiple public sources in order to systematically annotate the inferred networks.

The integrated transcriptomics analysis of public datasets shed light on a number of genes which are frequently deregulated in the psoriatic lesion compared with the unaffected skin in a large number of studies. In particular, *CRABP2, LCN2, S100A12* and *PDZK1IP1* were found to be deregulated in all of the datasets analyzed.

Furthermore, the analysis of co-expression networks highlights genes showing aberrant patterns of connectivity in the lesional network as compared to the network inferred from unaffected skin samples. For instance, we identified co-expression patterns of *SERPINB4, KYNU* and *S100A12* as being the most affected by the disease. Network analysis allowed us to identify *YPEL1* and *HUS1* as plausible, previously unknown, actors in the expression of the psoriasis phenotype. In addition, by exploiting topological properties of the network models, we highlighted a set of 250 non-deregulated genes, 223 of which have never been associated with the disease before, including *CACNA1A, HADH, ATP5MC1* and *CBARP* among others.

Finally, we characterized specific communities of co-expressed genes sustaining relevant molecular functions and specific immune cell types expression signatures playing a role in the psoriasis lesion. Overall, integrating experimental driven results with curated functional information from public repositories represents an efficient approach to empower knowledge generation about psoriasis and may be applicable to other complex diseases.

## Introduction

Psoriasis is a chronic inflammatory disorder of the skin, characterized by abnormal keratinocytes differentiation and hyper-proliferation of the epidermis, along with infiltration of inflammatory cells ^1^. Although genetic ^2^ and environmental factors are known to contribute to the aetiology of this polygenic disease, many of the intricate mechanisms of molecular alteration underlying the disease remain largely uncovered ^3^. Multiple transcriptome studies have pinpointed key pathways altered in lesional psoriasis skin ^4–8^. However, integrated analysis of multiple homogenized datasets is, to date, still limited to a few examples ^9,10^. Although, for instance, Piruzian and colleagues, ^9^ report the results of an integrated meta-analysis of both protein and gene expression datasets (and, therefore, integrating different data types), they still summarize the results of single datasets. Moreover, the analytical strategies employed in such studies have an impact on the ability to disentangle more complex patterns of molecular deregulation. In fact, while univariate differential expression analysis shed light on hundreds (sometimes thousands) of deregulated genes in the lesional skin, it is not straightforward to infer regulatory loops of molecular alterations underlying the phenotype.

This gap of knowledge could be filled by exploiting the large amount of biological data accumulated in recent years. In fact, vast amounts of data have been collected in public repositories and made freely available to the scientific community.

However, integrating such a wealth of data sources is still challenging due to the heterogeneity of data formats and the need for extensive manual curation ^11^.

A rigorous integration and exploitation of public data can provide a double benefit. On one side, already available data can inform the design of novel experimental strategies in order to achieve new knowledge. On the other hand, publicly available data may provide a shortcut to characterize and interpret *de novo* findings derived from targeted experiments. In the context of psoriasis, several repositories, among which Pharos (https://pharos.nih.gov/) ^12^, Target Validation [https://www.targetvalidation.org], Human Protein Atlas (https://www.proteinatlas.org) ^13^ and Clinical Trials (https://www.clinicaltrials.gov), report fundamental information about the state of the art of research in this topic, starting from the druggability/tractability of suitable drug targets to clinical trials and large scale genetic association studies. Such data has never been integrated in order to derive new knowledge about the mechanistic events underlying the psoriatic phenotype.

Graph theory provides effective models to uncover the relevant gene-gene expression relationships both in physiological and pathological conditions ^14,15^. In fact, gene coexpression network analysis is currently employed to understand the relationship between pairs of genes, and ultimately, gene networks or modules representing a marker of impaired biological functions in a disease ^16^.

In this study, we have integrated gene expression analysis and co-expression network analysis approaches on a large collection of manually curated transcriptomics datasets ^17^ in order to 1) validate and prioritize genes that are already known to be associated to psoriasis, 2) uncover novel genes never associated to psoriasis before, 3) create a gene-centric compendium of psoriasis-related information curated from multiple data repositories.

## Methods

### Data collection and preprocessing

All the raw transcriptomics data collected and utilized in this manuscript are publicly available in the Gene Expression Omnibus (GEO) repository. The preprocessed data consist of 23 microarray-derived gene expression datasets of both lesional (574 samples) and non-lesional skin (540 samples) of psoriasis patients. GEO IDs of the collected datasets are reported in Table S1.

The preprocessing procedure was carried out as described in Federico *et al*. ^17^. Differentially expressed genes for each dataset were identified through the use of the eUTOPIA software ^18^ by comparing the lesional skin samples with the non-lesional ones. For the analysis, eUTOPIA default parameters were used.

### Integrated large scale transcriptome analysis

The lists of differentially expressed genes were combined and the genes ranked on the basis of their frequency of differential expression across the datasets. Gene IDs conversions were performed through the use of the *bioMart* ^19^ and the *clusterProfiler* ^20^ Bioconductor packages. Similarly, for each list of differentially expressed genes, a functional annotation was performed by using the *ReactomePA* R package ^21^. The lists of significantly deregulated pathways were then combined and the pathways ranked on the basis of their frequency of deregulation across the datasets.

The gene and pathway rankings were carried out through the use of custom R scripts.

### Data scaling

All of the collected microarray datasets were combined for cross-platform normalization. In particular, the *pamr* R package (version *1.56.1*)^22^ was used to mean-adjust the combined microarray data based on a batch variable representing the different datasets downloaded from GEO.

### Integrated Psoriasis Knowledge Base construction

We built a comprehensive gene-centric annotation, namely Integrated Psoriasis Knowledge Base (IPKB), reporting aggregated information about psoriasis from several categories of databases. In detail, the IPKB contains information annotated in 14 databases, grouped in 6 categories: druggability/Tractability, Genetic association, Cell line-specific expression profiles, HumanKO/Trial, Immune Pathways and Modules, and literature derived PSO-association, for a total of 22 gene sets (Figure S2). The breakdown of the IPKB is reported in Table S2.

The IPKB was constructed by collecting data from numerous publications and or public databases. Available psoriasis genetic data were retrieved from the NHGRI-EBI GWAS catalog of published genome wide association studies ^23^ by using the keywords *“Psoriasis”* and *“Psoriasis vulgaris”* and selecting the genes with association p-value ≤ 1E-05, and the Open Targets database ^24^, selecting genes with genetic association score ≥ 0.1 for further analyses. Small molecule and antibody tractability data were also retrieved from Open Targets. Small molecule and biologics druggability data were collected from Finan et al., 2017 ^25^. Protein localization data were downloaded from Pharos^26^, Human Protein Atlas^27^ (URL: http://www.proteinatlas.org) and from Uva *et al*., 2010 ^28^. Immune pathway modules were retrieved from the Reactome database ^29^. Human knockout (KO) data were from Saleheen, *et al*., 2017 ^30^ and from Narasimhan, *et al*., ^31^. Immune cell-specific scRNA-Seq transcriptional signatures were collected from the Human Protein Atlas. The IPKB is publicly available in Zenodo (doi: 10.5281/zenodo.4740406).

### Networks inference and analysis

Two distinct co-expression networks were inferred by using the gene expression profiles of the lesional and non-lesional skin samples over all the included studies and the genes common to all the platforms. The co-expression networks were inferred through the use of the *INfORM* algorithm ^32^. We set up *INfORM* in order to build a robust consensus network by using the clr ^33^, aracne ^34^ and mrnet ^35^ algorithms with the following correlation and mutual information measures: Pearson correlation, Kendall correlation, Spearman correlation, empirical mutual information, Miller-Madow asymptotic bias corrected empirical estimator, Schurmann-Grassberger estimate of the entropy of a Dirichlet probability distribution and a shrinkage estimate of the entropy of a Dirichlet probability distribution, as implemented in the *minet* Bioconductor package ^36^. In order to carry out a network community detection we used the walktrap algorithm ^37^, implemented in *INfORM*. All computations performed on the inferred networks were carried out through the use of the *igraph* Bioconductor package ^38^.

### Functional annotation

The functional annotations carried out in this study were based on the Reactome biological pathways and performed through the use of the *ReactomePA* ^21^ and *clusterProfiler* Bioconductor packages ^20^. Moreover, the STRING database ^39^ was used to inspect the functional characteristics of the bridge genes.

### Visualisation

Visualisation of the results were performed through the use of the *ggplot2*^40^ and *gplots* ^41^ Bioconductor packages. The rendering of co-expression networks was performed by employing the *gephi* software ^42^. In this manuscript, we show a reduced representation of the actual networks in order to facilitate the reader visualisation.

### Differential centrality analysis

For each of the networks, their node betweenness, closeness and degree centralities were calculated with the Python’s *NetworkX* package (Python 3.6, NetworkX 2.3). The nodes were ranked according to each of the centrality measures. For each of the networks, their nodes’ median rank based on the rankings of the three centrality measures were calculated. To compare the network of the lesional skin with the non-lesional one, the absolute difference between the median ranks of the two networks was calculated and the genes were ranked accordingly.

### Gene set enrichment analysis

One tail gene set enrichment analyses (GSEA) were performed through Kolmogorov-Smirnov statistics, as implemented in the *stats* R package. Overrepresentation tests were performed by using the *bc3net* CRAN package ^43^.

#### druggability evaluation of the lesional network

The druggability evaluation of the PSO lesional network was performed by using the DrugBank annotation (version 5.1.7) ^44^. The Anatomical Therapeutic Chemical classification system (ATC codes) was retrieved from the *josetung/atc* Github R package. In order to increase the specificity of our analysis, we retrieved the drugtarget associations from DrugBank and considered only drugs whose targets are included in one module. The analysis was performed by considering the level 2 of the ATC codes annotation.

## Results

### *S100A12, PDZK1IP1, LCN2* and *CRABP2* are the most commonly upregulated genes in the psoriatic lesion

In order to identify genes that are consistently deregulated in transcriptomics experiments of lesional skin samples with respect to non-lesional counterparts, we first analyzed each dataset individually. The number of differentially expressed genes in each dataset ranged from 3,717 in GSE67853 to 100 in GSE57376, with a median of 1,863 (Supplementary Fig. S1). Therefore, we ranked the differential expressed genes on the basis of their occurrence across all the datasets. As a result, *S100A12, PDZK1IP1, LCN2*, and *CRABP2* genes were found to be differentially expressed in all 23 PSO datasets. Furthermore, a set of genes belonging to the *S100* and *SerpinB* transcription factor families was differentially expressed in 22 out of 23 datasets. Overall, 92 genes were differentially expressed in at least 20 datasets. The top 100 ranked gene list derived from the integrated gene expression analysis is reported in Supplementary Table S3.

We then assessed which genes showed the highest magnitude of deregulation across all the datasets. Therefore, we ranked each differentially expressed gene in each dataset by a significance score, calculated as follows:

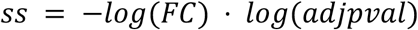

where *FC* is the fold change between the mean of the expression of the lesional samples and the mean of the expression of the non-lesional counterparts; *adjpval* is the Benjamini-Hochberg ^45^ adjusted p-value as obtained from the differential expression analysis. Our analysis highlighted *SERPINB4, S100A12*, and *TCN1* as the most deregulated genes over all the datasets (Fig. 1). Among the frequently upregulated genes, *SERPINB4* showed a median logFC across the datasets of 6.3 [Q1: 5.4; Q3: 7.2] with a maximum of 7.8 in GSE13355; *S100A12* showed a median logFC of 5.0 [Q1: 4.5; Q3: 6.0] and a maximum of 6.7 in GSE30768; and *TCN1* had a median value of 5.1 [Q1: 4.1; Q3: 5.4] and a maximum of 7.2 in GSE57376. On the other hand, the top genes found to be downregulated in most of the datasets were *BTC*, with a median logFC of −3.0 [Q1: −3.3; Q3: −2.6], and the strongest downregulation reported in GSE50790; *WIF1*, with a median logFC of −2.5 [Q1: −2.7; Q3: −2.3] and a maximum downregulation in GSE50790; and *PM20D1* with a median of 2,6 [Q1: −2.9; Q3: −2.0] and the lowest logFC of −4.5 in GSE47751.

**Figure 1.**
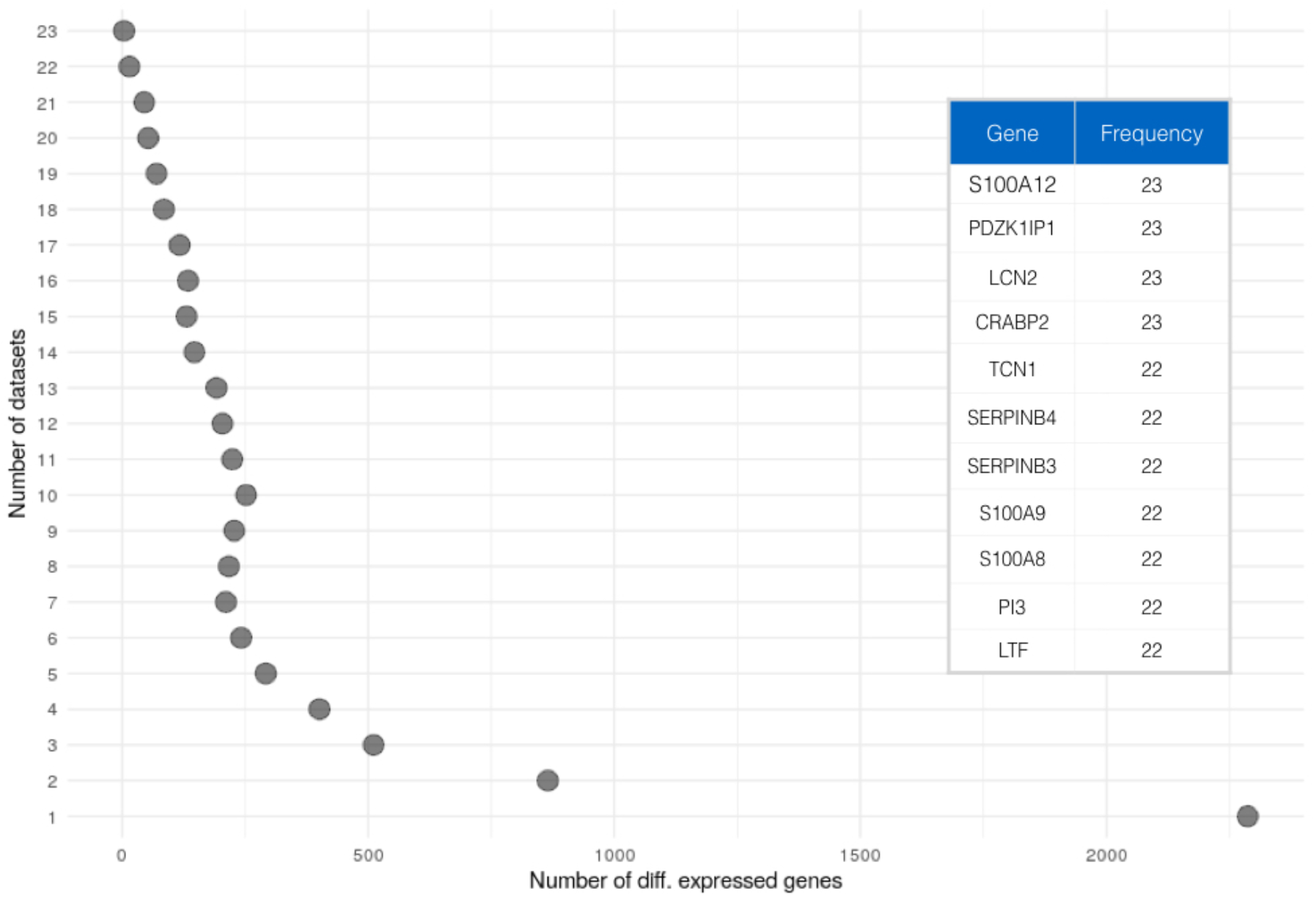
Occurrence of each gene as differentially expressed across the included studies (n=23). The table reports the top ranked genes and their differential expression frequency.

### Network analysis highlights genes with aberrant co-expression patterns

The integrated gene expression analysis allowed us to identify genes that are deregulated in the psoriatic lesion with respect to the non-lesional skin, giving, then, a quantitative perspective of the molecular alterations at a transcriptional level of the disease. However, the integrated expression analysis uncovers only one aspect of the deregulation underlying the psoriatic phenotype. In fact, the molecular build-up of a tissue is not only determined by the expression patterns of individual genes, but also by their co-expression relationships. Therefore, to characterize the complex landscape of transcriptional alterations that sustain the psoriasis, we identified disrupted patterns of gene co-expression. To do so, we inferred two transcriptome-wide gene coexpression networks from both the lesional and the non-lesional skin sample sets, respectively.

Since the networks were built from all the genes common to all of the microarray platforms, both of the networks are composed of 7,310 genes, while the lesional network has 1,136,431 edges, the non-lesional one has 1,559,790 edges.

The patterns of molecular alterations underlying the psoriatic phenotype can be characterized by investigating intrinsic topological properties of the inferred networks. One aspect that defines the differences between the two networks (lesional and non-lesional) is the centrality of their genes, a property measuring the number of coexpression connections that a certain gene holds with the others.

The differential centrality analysis allowed us to identify genes with a significant difference of connectivity between the lesional and non-lesional network. Table 1 shows genes whose connectivity is heavily affected by psoriasis, since they are highly central in the lesional network while their centrality is lower in the non-lesional one. Specifically, the connectivity of *SERPINB4, KYNU, S100A12, CASP5, CXCL1* and *CXCL8* are the most affected. Intriguingly, by comparing the results of the differential centrality analysis with the gene rank obtained by the integrated gene expression analysis, we observed that several of the top differentially central genes (DCG) were also differentially expressed in a large number of datasets. For instance, *SERPINB4, KYNU, S100A12, PNP* and *CXCL1*, which are among the top 10 DCG, resulted to be differentially expressed in more than 20 datasets. On the contrary, some genes such as *YPEL1* and *HUS1* appear at the top of the DCG but are not differentially expressed in any of the collected datasets.

**Table 1.** Top ten differentially central genes (DCG) between the lesional and the non-lesional network. The rank position in the lesional and non-lesional network, the difference between the ranks and the frequency of differential expression of each gene are reported.

| Top differentially central genes between lesional and non-lesional networks | Rank position in the lesional network | Rank position in the non-lesional network | Difference between the networks' ranks | Frequency of differential expression in the integrated expression analysis |
| --- | --- | --- | --- | --- |
| *SERPINB4* | 552 | 7,128 | 6,576 | 22 |
| *KYNU* | 911 | 7,046 | 6,135 | 22 |
| *S100A12* | 697 | 6,620 | 5,923 | 23 |
| *CASP5* | 1,043 | 6,875 | 5,832 | 11 |
| *CXCL1* | 1,107 | 6,218 | 5,783 | 20 |
| *CXCL8* | 1,781 | 7,135 | 5,354 | 18 |
| *SLC23A2* | 1,637 | 6,983 | 5,346 | 21 |
| *YPEL1* | 1,336 | 6,675 | 5,339 | 0 |
| *PNP* | 1,177 | 6,377 | 5,200 | 21 |
| *HUS1* | 1,048 | 6,232 | 5,184 | 0 |

On the other hand, we identified a second set of DCG, which showed an opposite pattern of aberrant connectivity compared to the genes reported in Table 1. Indeed, the connectivity of a number of genes is affected so that the genes are highly central in the non-lesional network while they show a lower centrality in the lesional one (Table 2). Therefore, these genes lose a high number of co-expression connections in the psoriatic lesion in respect of the uninvolved skin. *IHH, AQP9, ITGB8, CD55, CMA1* are the most affected ones, showing this trend of connectivity. Interestingly, their frequency of differential expression in the integrated expression analysis is markedly low, being detected as differentially expressed in a maximum of 2 datasets, with the exception of AQP9, detected in 17 studies. An overview of the impact of PSO on the co-expression connections in both the lesional and non-lesional network is shown in Supplementary Fig. S3.

**Table 2.** Top ten differentially central genes (DCG) between the lesional and the non-lesional network. The rank position in the lesional and non-lesional network, the difference between the ranks and the frequency of differential expression of each gene are reported.

| Top differentially central genes between lesional and non-lesional networks | Rank position in the lesional network | Rank position in the non-lesional network | Difference between the networks' ranks | Frequency of differential expression in the integrated expression analysis |
| --- | --- | --- | --- | --- |
| *IHH* | 6,575 | 1,447 | 5,128 | 0 |
| *AQP9* | 5,756 | 631 | 5,125 | 17 |
| *ITGB8* | 6,880 | 1,780 | 5,100 | 1 |
| *CD55* | 6,023 | 949 | 5,074 | 1 |
| *CMA1* | 5,874 | 852 | 5,022 | 2 |
| *SRPK3* | 6,711 | 1,693 | 5,018 | 0 |
| *ITPR1* | 6,458 | 1,529 | 4,929 | 1 |
| *TUBGCP3* | 5,865 | 971 | 4,894 | 0 |
| *SYCP2* | 6,570 | 1,820 | 4,750 | 0 |
| *NMB* | 6,389 | 1,726 | 4,663 | 3 |

### Identification of novel candidate genes associated with psoriasis

We hypothesized that, by studying the connectivity patterns among known psoriasis genes, it is possible to identify additional associated genes. Hence, a gene that is connected to two or more known psoriasis-associated genes is a strong candidate to be involved in its pathogenesis (Fig. 2). Based on this principle, we identified all the genes connecting pairs of differentially expressed genes (previously identified by the integrated gene expression analysis) within each of the networks (lesional and non-lesional, respectively), and hence acting as a bridge (hereafter referred to as “bridge genes”). By this analysis, we obtained a set of 1,622 and 1,940 bridge genes (BG) for the lesional network and non-lesional networks, respectively. Consequently, we selected a set of 250 genes acting as bridges in the lesional network, but not in the non-lesional one (Fig. 2). Among the bridge genes connecting a large number of differentially expressed gene pairs, we identified *CACNA1A (*Calcium Voltage-Gated Channel Subunit Alpha1 *A*) and its negative regulator *CBARP*, connecting 696 and 562 gene pairs, respectively. Likewise, the genes *HADH* and *ATP5MC1*, whose protein products function in mitochondria, connect a high number of deregulated gene pairs (562 and 550, respectively) in the lesional network.

**Figure 2.**
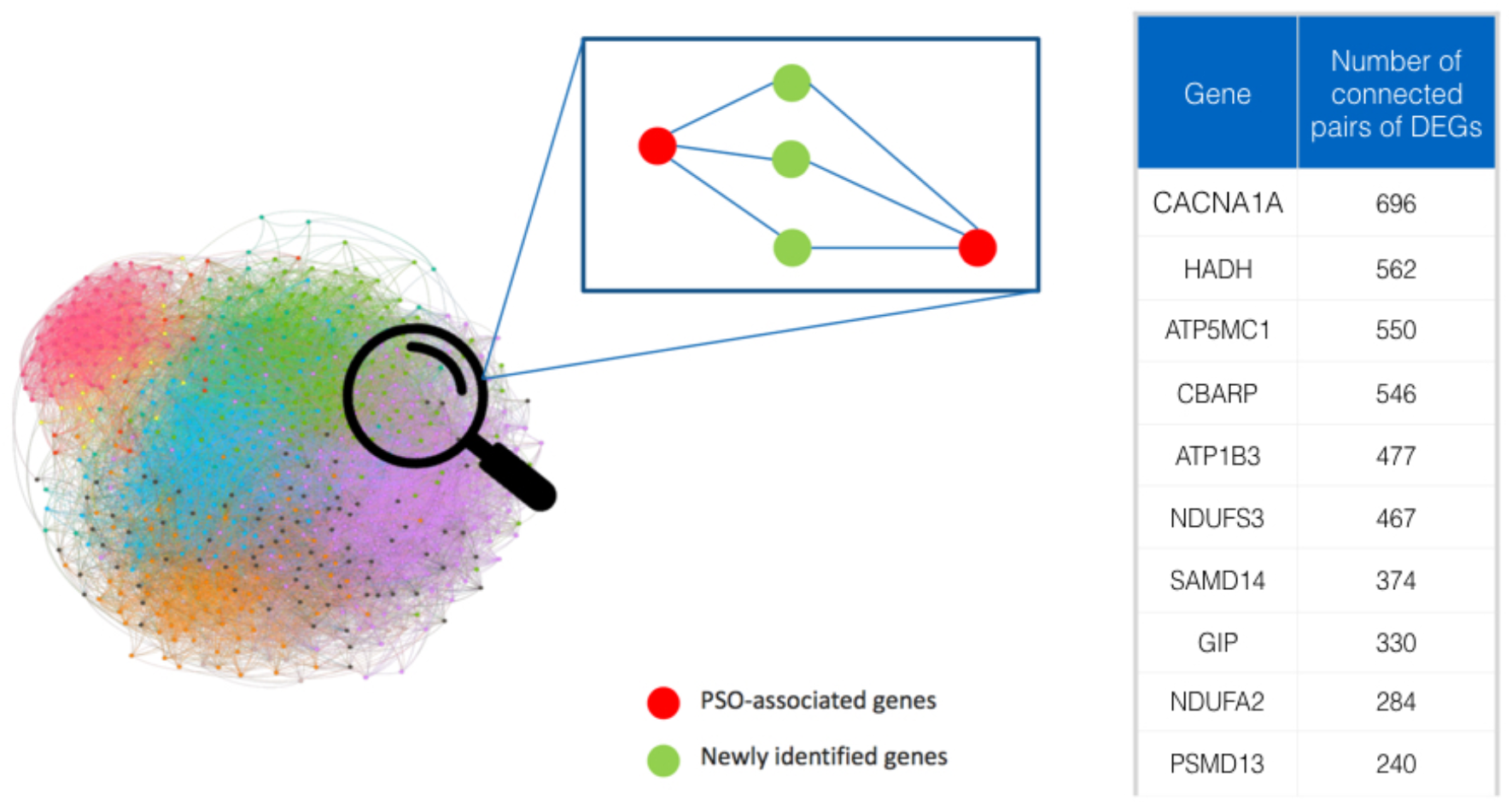
Schematic representation of bridge genes. In blue is shown as an example of gene co-expression network. In green are shown the bridge genes, acting as connectors among couples of differentially expressed genes, shown in red. The table reports the rank of bridge genes based on the number of connected couples of differentially expressed genes.

In order to characterize the functional properties of the bridge genes, we performed a functional annotation by using the STRING database. STRING shows a peculiar clustering of gene products involved in *RNA splicing*, which is the first enriched term in the gene ontology (GO) biological process, followed by cellular *nitrogen compound metabolic process* (both with FDR = 0.0063) (Fig. S4).

### Network analysis allows the identification of disease-relevant communities

It is a widespread assumption that genes which are tightly co-expressed (whose expression levels are highly correlated) are likely to be also co-regulated, as well as involved in common functions ^46^. Graph models allow the identification and characterization of such communities of genes. In this study, we investigated the arrangement of co-expressed genes in both the lesional and non-lesional networks by performing a community detection analysis. Thus, we identified 13 communities of coexpressed genes in the lesional network. The biggest community encompasses 1,888 genes, while the smallest 1 gene, with a median size of 309. In the non-lesional network we identified 10 communities with a median size of 756 genes, being the biggest composed by 1,723 and the smallest by 1 gene. All our analyses were limited to modules composed by at least 10 genes (Supplementary Fig. S5 and S6).

An interesting aspect we investigated is whether one or more network communities exist that enrich for the putative PSO-associated genes identified through the integrated gene expression analysis. In order to fulfill this aim, we performed a GSEA on the gene rank derived from the integrated gene expression analysis over the identified modules of the lesional network. As a result, we obtained that module 2, module 4 and module 7 significantly enriched for the genes at the top of the integrated gene expression analysis rank (*p*=2.41e-23, *p*=1.25e-24, *p*=4.89e-07, respectively). Similarly, we performed the same analysis to assess whether the communities of the lesional network enrich for genes whose centrality is significantly different between the lesional and the non-lesional network, previously identified by the differential centrality analysis. We obtained that module 6 and module 7 significantly enrich for the top differentially central genes (*p*=0.0054 and *p*=0.0026, respectively). Finally, the same analysis was performed on the bridge genes set, in order to verify their enrichment over the modules. We found that module 3 significantly enriches for the bridge genes (*p*= 1.57e-07) (Fig. 3).

**Figure 3.**
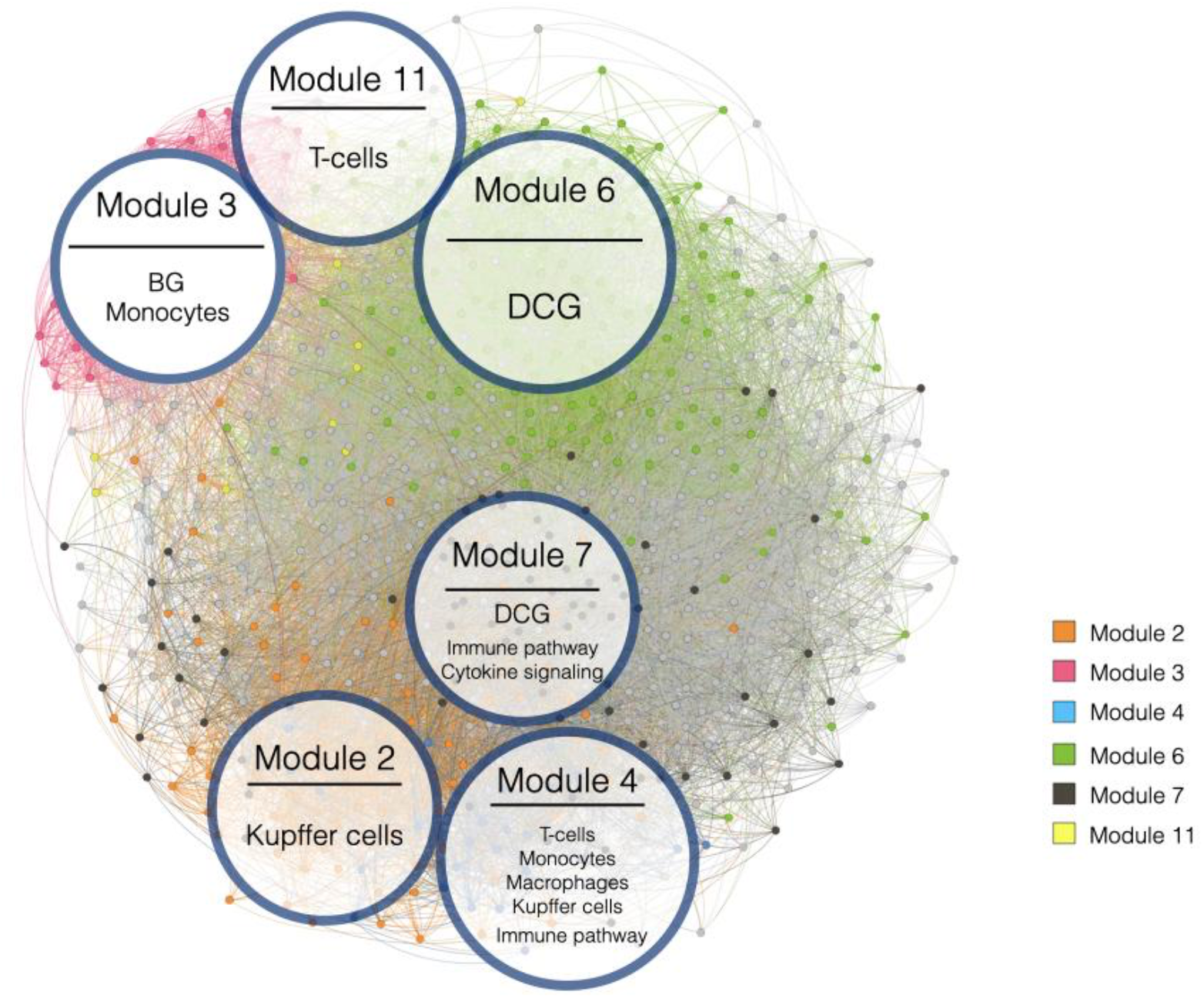
Evaluation of the enrichment of differentially central genes (DCG), bridge genes (BG), immune cell specific genes as well as immune-related pathways over the modules detected in the lesional network. The size of the circle is proportional to the dimension of the module.

We characterized the biological functions of the gene communities identified in the lesional network (*fdr* < 0.05, Fig. 4). Module 2, which is overrepresented by top-ranked genes of the integrated gene expression analysis, is significantly enriched in genes belonging to the extracellular matrix organization and proteoglycans, *collagen formation*, *integrin cell surface interactions* among others, which are expected in skin diseases like psoriasis. *Interleukin 4 and interleukin 13 signaling* pathways are also enriched by module 2 and 7. Moreover, *interleukins signaling* pathway is also enriched in module 7, together with other immunological pathways, such as *interferon signaling* pathway. However, Module 4 shows the strongest immunological signature among all. In fact, the genes belonging to this module significantly enrich *interleukin 10 signaling, interleukin 4 and 10 signaling* and *interferon alpha/beta signaling*. Module 5 and 6 overrepresent pathways related to generic cell cycle functions, like *G1/S transition, S phase, transcriptional regulation of P53, mRNA splicing*. Likewise, module 3, which is over-represented by bridge genes, enriches mostly for receptorial functions, such as *G-Protein Coupled Receptors (GPCRs) ligand binding, rhodopsin-like receptors*, and *peptide ligand-binding receptors*.

**Figure 4.**
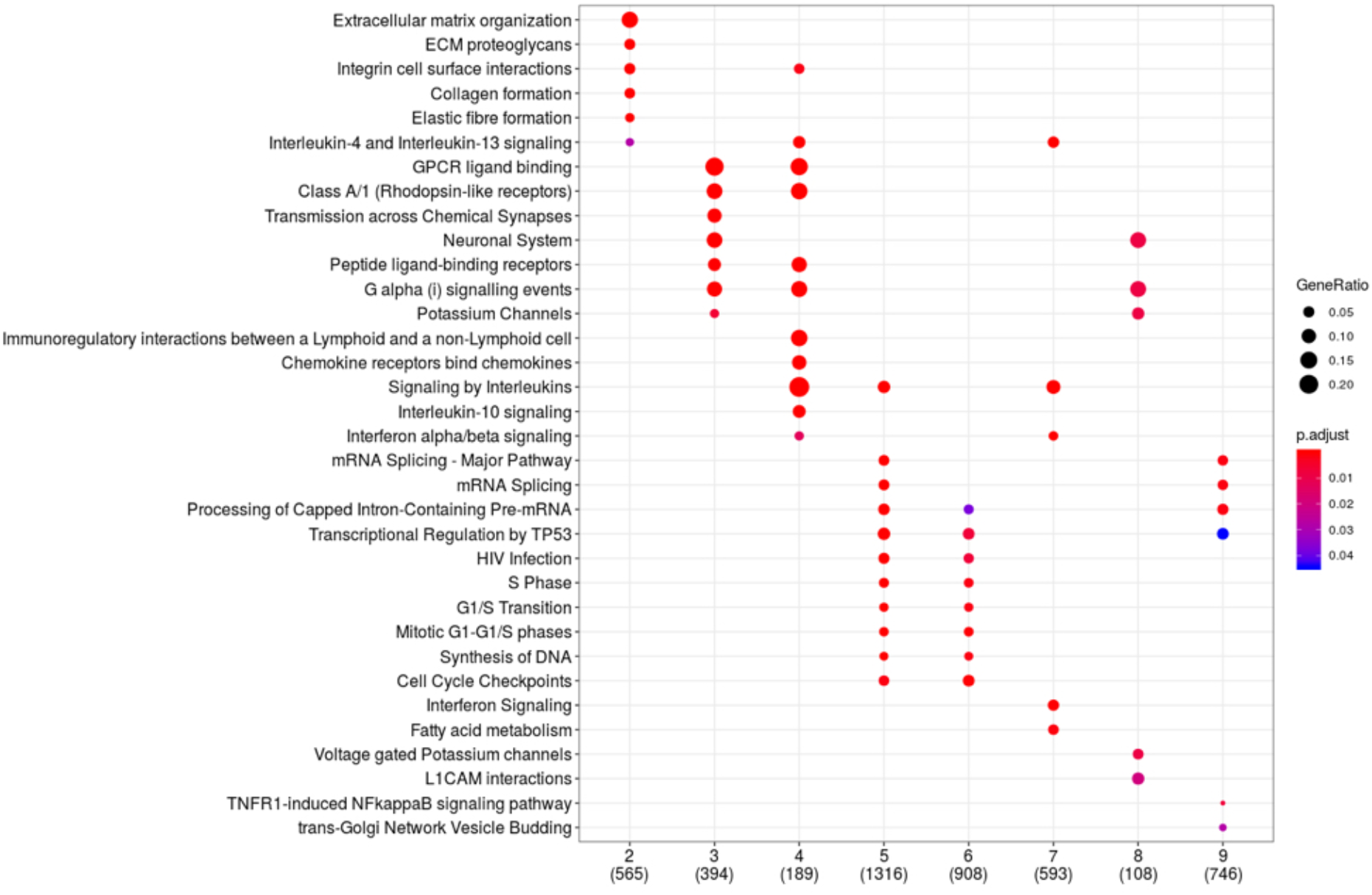
Module-specific pathway enrichment based on the Reactome database. On the x axis are indicated the module and the number of genes contributing to the enrichment (in parentheses). On the y axis are indicated significantly enriched pathways.

### Prior knowledge enables the characterisation of lesional network gene modules

We further performed an overrepresentation analysis of the gene sets collected in the IPKB in each community of the lesional network. This analysis highlighted that, for the category Human KO/Trial, Module 4 is significantly enriched in two out of three gene sets, HumanKOPakistan and ClinicalTrial (*p*=0.027 and *p*=0.001, respectively). Module 3 and 7 enrich for genes belonging to the HumanKOPakistan and HumanKOBritishPakistani sets (*p*=0.003 and *p*=0.001, respectively). Module 4 is also enriched for almost all the gene sets of the category Immune Pathways/Modules, with the most significant *p*=5.06E-18 in ImmunePathwayAdaptive. On the other hand, Module 7 enriches for the ImmunePathwayCytokineSignalling (*p*=2.12E-05). Additionally, since psoriasis poses its roots in the impairment of the immuno-inflammatory homeostasis, we wondered whether the modules of the lesional network are enriched by genes expressed in a specific manner in immune cell lines, which are primarily involved in the aberrant response in psoriasis. By exploiting publicly available immune cell type-specific gene expression signatures from the Human Protein Atlas database, which we included in the DGI, we performed a GSEA analysis to assess the enrichment of cell type-specific genes over the modules of the lesional network. We obtained that module 4 is enriched by genes specifically expressed in T-cells (*p*=0.006), monocytes (*p*=7.93e-05), macrophages (*p*=0.015) and Kupffer cells (*p*=0.027). Similarly, module 2 is enriched by monocytes-specific genes (*p*=0.035) while module 3 by Kupffer cells genes (*p*=0.035) (Fig. 3).

### Immunomodulators and dermatological drugs target specific modules of the lesional skin network

We characterised the druggability potential of the relevant modules identified in the previous analytical steps. To this end, we defined module-specific drug-target gene maps by exploiting publicly available information available at DrugBank. All of the modules except module 5 and 9, encompass a number of drugs which is higher than the number of genes composing the module (Fig. 5). Moreover, by considering modules with a number of genes >10 and taking into account the frequency of druggable genes over the total number of genes composing each module we found that module 3, 8, 7, 2 and 4 show a higher amount of drug target genes (38%, 38%, 31%, 31%, 30%, respectively) compared to other modules.

**Figure 5.**
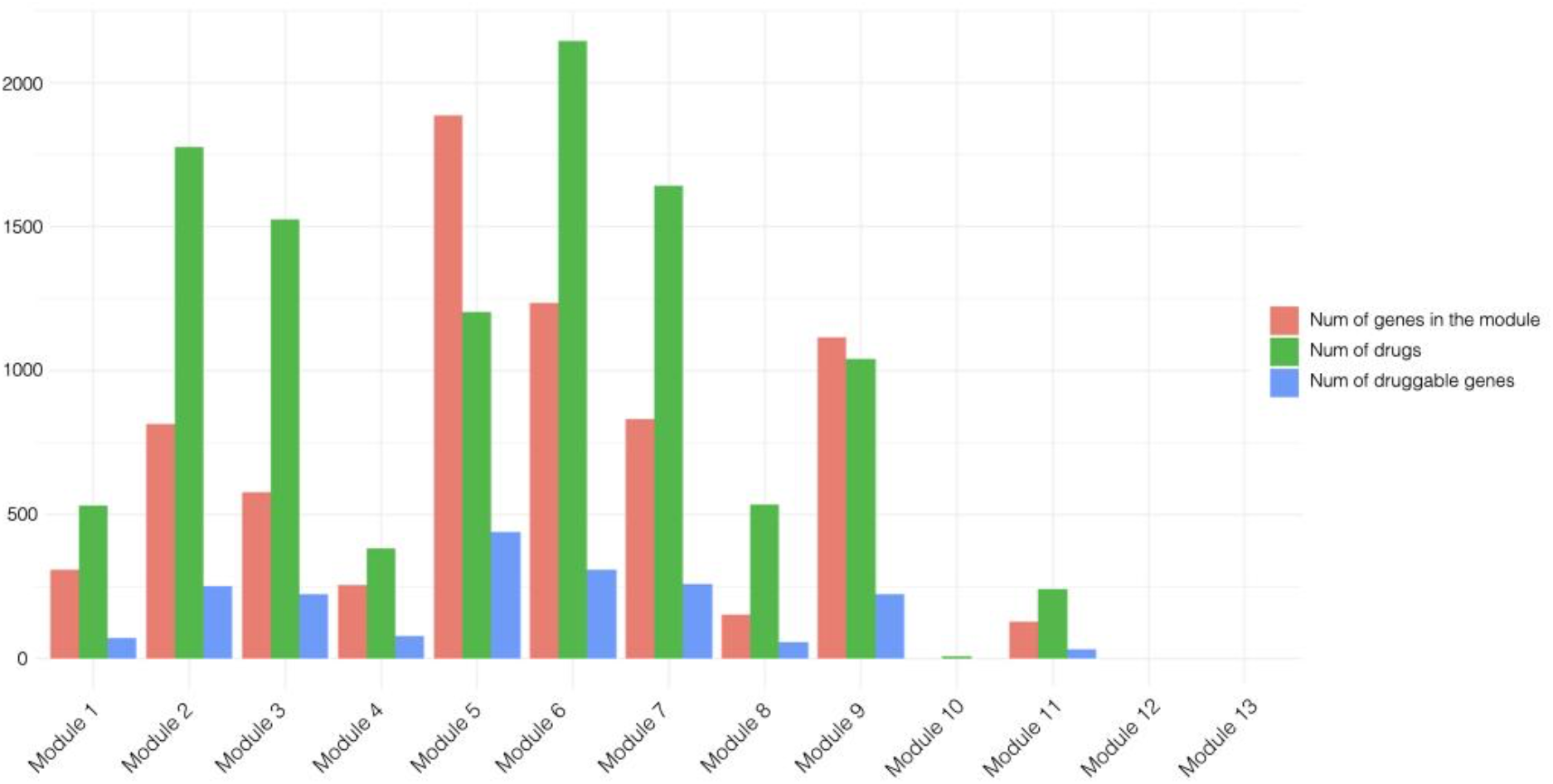
Evaluation of the module-based druggability profile in the PSO lesional network. In brown is shown the number of genes composing the module, in green the number of drugs and in blue the number of druggable genes.

To further characterise the drugs mapping onto the lesional network, we restricted our analysis to drugs whose targets belong to one specific module only. By applying this restraint, we carried out further analyses on 3090 out of 5317 drugs.

We characterised the module-specific drugs on the basis of their therapeutic class as annotated in the second level of the World Health Organization (WHO) Anatomical Therapeutic Chemical (ATC) classification system. Module 8 contains the highest number of drugs mapped with respect to the size of the module (140 drugs, drugs/genes ratio = 0.92), followed by module 11 (87 drugs, drugs/genes ratio = 0.68), module 2 (500 drugs, drugs/genes ratio = 0.61), module 3 (354 drugs, drugs/genes ratio = 0.61), module 6 (729 drugs, drugs/genes ratio = 0.59). Module 4, which we previously identified to have a marked immunological profile, encompasses target genes for 106 drugs, showing a drugs/genes ratio of 0.41.

While the most represented drug category in module 8 is anti-emetics and anti-nauseants (A04), for both module 8 and module 11 dermatologicals belonging to the anti-acne preparations category (D10) are predominantly represented (Fig. 6). Potassium Voltage-Gated Channel Subfamily H Member 2 (*KCNH2*) and the Retinoic Acid Receptor Alpha (*RARA*) play a pivotal role in the druggability of module 8. In fact, KCNH2 protein is a target of a high number of drugs, including Erythromycin and Chlorobutanol, while the retinoic acid receptor alpha is targeted by dermatological compounds including tretinoin, isotretinoin and adapalene. Interestingly, *RARA* is also targeted by two other retinoids employed in the treatment of severe psoriasis, Tazarotene and Etretinate. In module 11, Dapsone and Resorcinol target the *NAT2* and *TPO* gene products, respectively. In module 4, the most represented class of compounds is immunosuppressant (L04). In fact, Alefacept, targeting the T-cell surface antigen CD2, together with Abatacept, Belatacept, targeting the T-lymphocytes activation antigen CD86 are among the numerous compounds belonging to this category. In module 4, also Framycetin is represented in the medicated dressings category (D09), which is known to act on the *CXCR4* gene product. Interestingly, module 4 also encompasses a number of target genes for immunostimulant compounds (L03). Our analysis highlighted that *IL2RA* and *IL2RB* are targets of Aldesleukin, a compound employed in *IL2* replacement therapies, while the Colony Stimulating Factor 3 Receptor (*CSF3R*) is targeted by several immunostimulant drugs, such as Filgrastim, Lenograstim, Pegfilgrastim, Lipegfilgrastim. Finally, module 2 is enriched by a wide spectrum of pharmacological categories, ranging from drugs employed in the treatment of musculo-skeletrical disorders (M09), hematological agents (B06) to drugs used for gastrointestinal disorders (A03) and anti-Parkinson drugs (N06).

**Figure 6.**
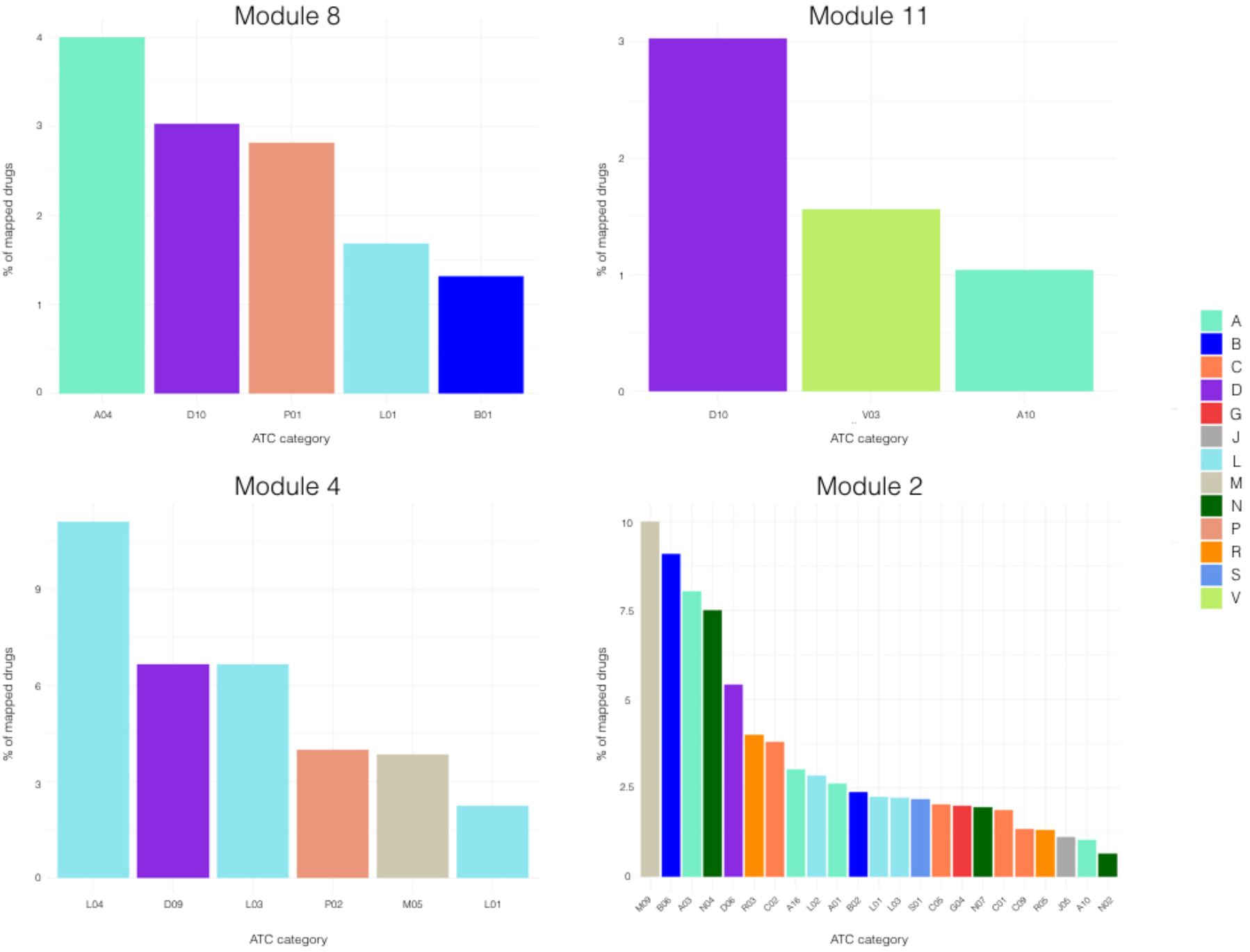
Characterisation of module-specific drugs based on the Anatomical Therapeutic Chemical classification system (ATC). The figure shows the modules with the highest drugs/genes ratio. The plots of the remaining modules are shown in Supplementary materials.

## Discussion

In this study, we analysed a large collection of transcriptomics datasets recently curated ^17^ in order to gain new knowledge about complex patterns of gene alteration with a role in the psoriatic phenotype.

By analyzing 23 transcriptomics datasets, we identified genes and pathways that are consistently deregulated in the psoriatic lesions as compared to the uninvolved skin. Our analysis found *CRABP2, LCN2, S100A12* and *PDZK1IP1* deregulated in all the datasets, suggesting their importance in the definition of the psoriatic phenotype. Overall, our differential expression analysis highlighted the upregulation of genes involved in inflammatory cascades, such as *S100A12* and *SERPINB4*, and the pronounced downregulation of genes related to developmental pathways, such as epidermal growth factor family members (*BTC*) and genes involved in the WNT signalling (*WIF1*).

In addition to known genes associated with psoriasis, we also prioritized novel candidates, such as *SYNCRIP* encoding a protein involved in the control of translation such as alternative splicing and mRNA maturation ^47^, as well as *SASH3* whose protein product could function as a signaling adapter proteins in lymphocytes ^48^.

Since it is well known that genes act in a coordinated manner in both physiological and pathological conditions, we inferred and analyzed co-expression network models representative of the psoriatic lesion and the uninvolved skin in order to identify the disrupted patterns of gene co-expression underlying the psoriatic lesion.

In the context of graph models, genes are co-expressed with variable numbers of other genes (interactors), signifying their relative importance in defining the phenotype underlying the gene network. We identified the genes with the most different number of interactors in the two networks derived from lesional and nonlesional samples, respectively.

This result highlights two important aspects. First, the deregulation of distinct genes in the lesional skin affects the co-expression relationships with other genes, which are not necessarily deregulated. Second, this highlights the importance of going beyond the classical gene expression analysis, which is focused on the evaluation of individual genes, failing to capture the complex relationships sustaining biological processes. In fact, we identified a few genes with an aberrant co-expression connectivity in the lesional network as compared with the non-lesional one, which do not show a differential expression. Among others, *YPEL1* and *HUS1*. While *YPEL1* may play a role in the regulation of cell division and in the polarization of fibroblasts towards an epithelial-like morphology ^49^, *HUS1* product is involved in the arrest of cell cycle upon DNA damage. In fact, *HUS1* gene is one of the top associated genes with Xeroderma pigmentosum variant, since it is responsible for the impaired repair capacity of UV-mediated DNA damages.

While differential expression analysis identified genes deregulated in psoriatic lesions, gene network analysis investigating the connectivity patterns among such genes in the lesional network revealed 250 non-deregulated genes connecting (bridging) deregulated ones. As previous attempts to identify genes associated with psoriasis by transcriptomics relied mainly on differential expression, it is not surprising that 223 of our 250 newly identified genes have not been associated with psoriasis so far according to Opentargets. Thus, we here describe a completely new group of genes related to transcriptional deregulation in psoriatic lesions, which we call “bridge genes”, since they connect couples of differentially expressed genes within the lesional network, and, therefore, they are putatively associated to psoriasis.

The bridge gene connecting the highest number of differentially expressed genes is *CACNA1A* (Calcium Voltage-Gated Channel Subunit Alpha1 A). This gene encodes a calcium channel, which regulates intracellular processes such as contraction, secretion, neurotransmission and gene expression, suggesting that bridge genes have superior/broad-spectrum roles in cell regulation. *CACNA1A* not only is followed by its related gene *CBARP* (CACN Subunit Beta Associated Regulatory Protein), but also by a number of genes involved in mitochondrial metabolic activities such as *HADH,* whose enzymatic activity is exploited in the fatty-acid beta-oxidation process, and *ATP5MC1*, coding for a subunit of the mitochondrial ATP synthase and responsible for the synthesis of ATP during oxidative phosphorylation by exploiting the protonic gradient across the mitochondrial inner membrane.

The analysis of communities of co-expressed genes allowed us to both identify genes that can be functionally involved in psoriasis and their characteristics in terms of immune cell-specific expression and druggability.

Interestingly, we found that many bridge genes are significantly co-expressed within module 3, which is enriched by genes involved in biological processes such as GPCR ligand binding, transmission across chemical synapses, and potassium channels indicating that bridge genes are related to broadly receptorial functions.

Pilar Pedro *et al*. ^50^ reports about the role of the GPCRs in the translation of extracellular signals into intracellular cascades that regulate the activation of keratinocytes proliferation and differentiation, including major signalling pathways, such as Hedgehog, Hippo YAP1 and WNT/B-catenin. In the same work, the authors underline the role of the neural-epithelial connection, mediated by β-adrenergic receptor (βAR) signaling in triggering keratinocyte proliferation, which is over-activated in the psoriatic lesion.

Moreover, module 3 is particularly rich in genes which are expressed in a specific manner in monocytes. The role of hyper-reactive monocytes in the psoriatic phenotype has long been known ^51^. In fact, Golden *et al*., observed elevated adhesion of monocytes and, in turn, increased formation of aggregates, which they also correlated with disease severity and underlying a major role of the innate immunity in the disease progression ^52^.

Module 4 is overrepresented by deregulated genes whose activity lies in immune-related pathways, including signaling by interleukins, interferon signaling and chemokines and their receptors. Moreover, it encompasses genes which are specifically expressed in a reservoir of immune cell lines, such as T-cells, monocytes, macrophages and Kupffer cells, underlying its role in the chronic auto-inflammatory response characteristic of psoriasis. Indeed, a pivotal role for T-cells, and cells of the myeloid lineage, including monocytes and macrophages is well established ^53–58^.

The immune-related nature of module 4 is reflected also by the druggability analysis of the lesional network model. In fact, several genes belonging to module 4 are targets of both immunostimulant and immunosuppressive drugs, such as interleukins and chemokines. This suggests that this module could be a good reservoir of putatively novel pharmacological targets for the development of therapeutic approaches with an immunomodulatory action to treat psoriatic lesions. Along with these categories of compounds, dermatological medications were also represented in module 4. Framycetin (also known as neomycin sulphate), among others, is a neomycin component employed in the treatment of ocular and skin bacterial infections. To the best of our knowledge, this compound is currently not employed for the treatment of the psoriatic plaques. Furthermore, module 8 showed an interesting *scenario* regarding its drug target content. In fact, we found that the retinoic acid receptor alpha (*RARA*), is the target of a plethora of chemical compounds already employed in the treatment of severe psoriasis plaques. For instance, the topical agent Tazarotene, and oral agent Acitretin (and its predecessor Etretinate), are compounds largely used in the treatment of psoriatic plaques ^59,60^. Tazarotene is a retinoid drug which has been approved in 2019 from Food and Drug Administration (FDA) in combination with Halobetasol in PSO affected adults^61^. On the other hand, Acitretin is used in severe psoriatic manifestations, but due to its high lipophilic capacity shows teratogenic effects and it is contra-indicated in pregnancy and for 3 years prior to conception ^62^. Etretinate, a metabolic product of Acitretin, is a high lipophilic retinoid which was used in severe psoriatic manifestations ^63^, but its use was suspended between 1996 and 1998 for its teratogenic effects ^64^.

The limited amount of clinical data made available along with the transcriptional profiles annotated in public repositories poses some limitations to the present study. The lack of detailed clinical information makes the identification of gene markers or co-expression communities associated with clinical characteristics impossible, hampering the predictive power of the present study. Moreover, this hinders the possibility of translating this study to a precision medicine level, making possible the characterization of the impaired molecular relationships at a single-patient resolution.

In conclusion, in this study we combined an integrative gene expression analysis with co-expression network analysis in order to identify novel aspects of the psoriatic lesion at a molecular level. Our approach allowed us to give an insight into the known alterations associated with psoriasis by identifying novel genes which can putatively act as disease biomarkers. Future mechanistic studies elucidate their role in the disease onset and progression while epidemiological studies will be necessary to assess their clinical relevance.

## Acknowledgements

This study was supported by the EU IMI2 Biomap Project (Grant agreement 82151).

## Author contributions

AF and DG designed the study. AF performed the analyses. AP and VF contributed to the analyses. DMK and EDR participated in data generation and curation. AF, LM, GdG, CS, SW and DG wrote the manuscript. DG supervised the study.

